# Theoretical analysis of the distribution of isolated particles in the TASEP: Application to mRNA translation rate estimation

**DOI:** 10.1101/147017

**Authors:** Khanh Dao Duc, Zain H. Saleem, Yun S. Song

**Author notes:** Also at Chan Zuckerberg Biohub, San Francisco, CA 94158.

## Abstract

The Totally Asymmetric Exclusion Process (TASEP) is a classical stochastic model for describing the transport of interacting particles, such as ribosomes moving along the mRNA during translation. Although this model has been widely studied in the past, the extent of collision between particles and the average distance between a particle to its nearest neighbor have not been quantified explicitly. We provide here a theoretical analysis of such quantities via the distribution of isolated particles. In the classical form of the model in which each particle occupies only a single site, we obtain an exact analytic solution using the Matrix Ansatz. We then employ a refined mean field approach to extend the analysis to a generalized TASEP with particles of an arbitrary size. Our theoretical study has direct applications in mRNA translation and the interpretation of experimental ribosome profiling data. In particular, our analysis of data from *S. cerevisiae* suggests a potential bias against the detection of nearby ribosomes with gap distance less than ~ 3 codons, which leads to some ambiguity in estimating the initiation rate and protein production flux for a substantial fraction of genes. Despite such ambiguity, however, we demonstrate theoretically that the interference rate associated with collisions can be robustly estimated, and show that approximately 1% of the translating ribosomes get obstructed.

## INTRODUCTION

The Totally Asymmetric Exclusion Process (TASEP) is a classical stochastic model for transport phenomena in a non-equilibrium particle system. Although it has been widely studied by mathematicians and physicists, the TASEP was first introduced in a biological context by McDonald et al. [1] to model mRNA translation and describe the dynamics of ribosomes moving along the mRNA. Over the past fifteen years, the TASEP and its extensions have been used for this purpose [2–11], and TASEP-based models have been recently applied to infer the translation rate from experimental measurements [9], in particular ribosome profiling data [12–14]. Ribosome profiling (also known as Ribo-Seq) is an experimental technique developed to examine position-specific densities of ribosomes along each mRNA [15], and thus captures the dynamics of mRNA translation to some extent. However, analytical tools for interpreting ribosome profiling data are still much in need of development [16], as relating the observed footprint density to the corresponding protein production rate remains challenging for several reasons [17]. One notable issue comes from the experimental protocol used to generate the ribosome profile. In general, long mRNA fragments that may account for stacked ribosomes are not sequenced. As a result, the observed density may only include well-isolated ribosomes, thus leading to a bias that needs to be corrected when evaluating the ribosome density [6, 14, 17–19]. Although the TASEP has been broadly studied under different conditions and using various approaches [20, 21], to our knowledge, the density of isolated particles has not been studied previously.

These theoretical and technical issues motivate us to study the extent of isolated particles in the TASEP, in order to quantify the relation between the mRNA translation dynamics and the observed densities in ribosome profiling data. To do so, we first employ the matrix formulation of Derrida *et al.* [22] to derive exact formulas for the density of isolated particles in the classical TASEP model, in which each particle is pointlike and occupies a single site. For the case when the number *N* of sites is large, we obtain simple asymptotic formulas. We then extend our study to the general case with particles of an arbitrary size. Using a refined mean field approach introduced by Lakatos and Chou [2], we derive new asymptotic formulas that agree well with Monte Carlo simulations.

We obtain new results regarding the translation dynamics by applying our theory to ribosome profiling data. In particular, our analysis of undetected ribosomes suggests a potential bias against consecutive ribosomes less than ~3 codons apart. Using a measurement of ribosome density called “translation efficiency” (TE), we provide estimates of the interference rate associated with traffic collision, and find that, for a significant fraction of genes, there is some ambiguity in identifying the initiation rate and the flux from TE. Although the TE has been widely used as a proxy for protein production rate [23], these results suggest that more refined methods and estimates should be used to properly quantify gene expression at the translation level.

## THEORETICAL RESULTS

In this section, we briefly introduce the TASEP model, and present our main theoretical results on the classical and generalized versions of model. Appendices A and B summarize some previously known results used in our analysis.

### A. The density of isolated particles in the classical TASEP model

We first studied the density of isolated particles in the context of the classical TASEP model with open boundaries [24]. Briefly, the dynamics of this stochastic process can be described as follows (see Fig. 1a). On a onedimensional lattice of *N* sites, the classical TASEP describes the configuration of pointlike particles, described by a vector ***τ*** = (*τ*_1_,…, *τ_N_*) such that *τ_i_* = 0 if the *i*th site is empty and *τ_i_* = 1 if it is occupied. During every infinitesimal time interval *dt*, each particle at site *i* ∈ 1,…, *N* − 1 has probability *dt* of hopping to the next site to its right, provided that the site is empty. Additionally, a new particle enters site 1 with probability *αdt* if *τ*_1_ = 0. If *τ_N_* = 1, the particle at site *N* exits the lattice with probability *βdt*. The parameters *α* and *β* are respectively called the initiation and termination rates. In the long time limit, the system reaches steady state and the corresponding expected marginal density of particles at position *i* on a lattice of size *N*, denoted 〈*τ_i_*〉_*N*_, is defined as,

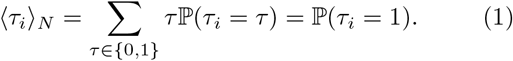

**FIG. 1.**
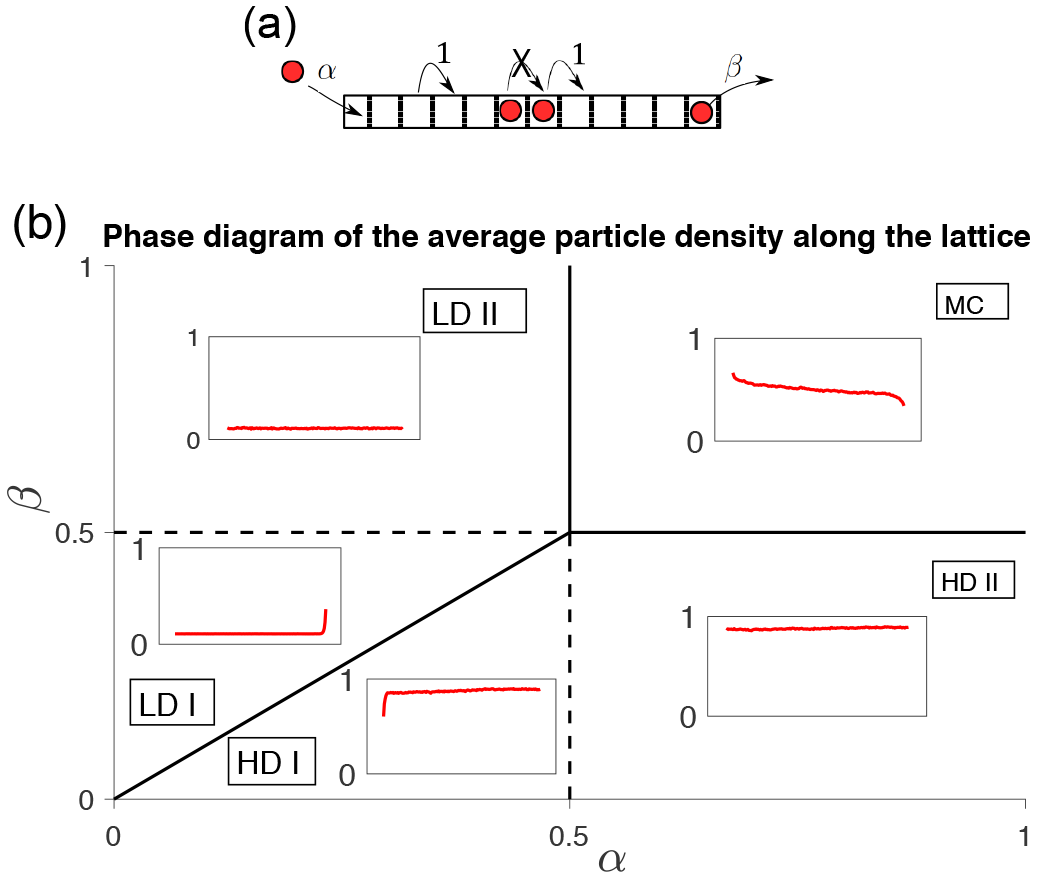
Illustration of the TASEP with open boundaries. **(a):** A schematic representation of the TASEP model. Particles are introduced at the start of the lattice with exponential rate *α* and move along with exponential rate 1, provided that there is no particle occupying the next site. At the end of the lattice, they exit with exponential rate *β*. **(b):** Phase diagram of the average particle density along the lattice. The profile of average density of particles along the lattice can be classified according to a phase diagram in (*α, β*)-space, separating different regions: the maximal current regime (MC), the low density regime (LD), and the high density regime (HD). The LD and HD regions can also be decomposed into two separate ones: LD I/II, and HD I/II, respectively.

Averaging the process over the events that may occur between *t* and *t* + *dt* leads to a system of equations relating one-point correlators to two-point correlators [25]. Similarly, two-point correlators can be related to three-point correlators (see Appendix A), and so on. To derive analytic expressions for the average densities, Derrida *et al.* [22] showed that the steady state probability of a given configuration can be derived using a matrix formulation satisfying a set of algebraic rules (see Appendix B). Using these rules, they obtained an exact formula for 〈*τ_i_〉_N_*, and showed that, in the large-*N* limit, the TASEP follows different dynamics according to a phase diagram in (*α, β*)-space.

In our work, we employed the aforementioned matrix formulation to derive analytic expressions for the average density of isolated particles. Specifically, consider the random variable 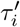 defined as

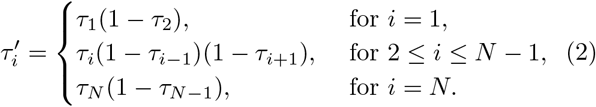

Note that 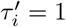 if there is an isolated particle at position *i*, and 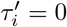 otherwise. From (2), we see that the average density 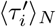 of isolated particles at an interior site *i*, where 2 ≤ *i* ≤ *N* − 1, is given by

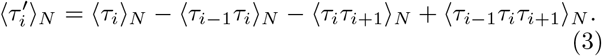

As detailed in Appendix C, by analyzing the terms on the right hand side of (3), we obtained, for 2 ≤ *i* ≤ *N* −1,

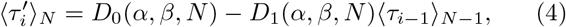

where

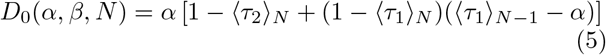

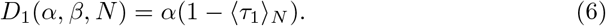

For the boundaries, we obtained

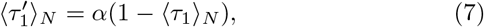

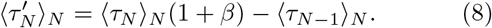

As mentioned earlier, exact formulas for 〈*τ*_*i*_〉_*N*_ are known [22] (see Appendix B), so plugging them into (4)–(8) leads to exact results for the average densities of isolated particles along the lattice.

### B. Large-*N* asymptotics in three different phases

We next derived the large-*N* asymptotics of 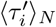 from those of 〈*τ_i_*〉*_N_*. In this section, we drop the dependence on *N* and write 〈*τ_i_*〉 instead of 〈*τ_i_*〉*_N_*. In the large-*N* limit, the dynamics of the TASEP can be separated into three different phases—namely, maximal current (MC), low density (LD), and high density (HD)—depending on the values of (*α, β*) (see Fig. 1b and (9) below). At steady state, 〈*τ_i_*(1 − *τ*_*i*+1_)〉 is the same for all *i* = 1,…, *N* − 1. This quantity is defined as the current (or flux) and is denoted by *J*. Using the asymptotics of the particle densities in the three phases [22], we found that *D*_0_(*α, β, N*) and *D*_1_(*α, β, N*) in (5) and (6), respectively, are both asymptotically equivalent to the asymptotics of *J* in the large-*N* limit, given by

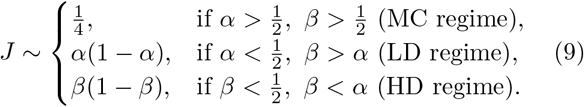

Hence, it turns out that the asymptotics of 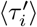 for 2 ≤ *i* ≤ *N* − 1 are correctly given by using in (3) the mean-field approximation 〈*τ*_*i*−1_*τ*_*i*_(1 − *τ*_*i*+1_)〉 ~ 〈*τ*_*i*−1_〉〈*τ*_*i*_(1−*τ*_*i*+1_)〉 = *J*〈*τ*_*i*−1_〉. Finally, noting that 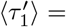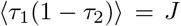 and *β*〈*τ*_*N*_〉 = *J* at steady state, while 〈*τ*_*N*−1_〉 ~ *J* + (*J/β*)^2^ asymptotically, we obtain that 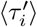 is asymptotically given by

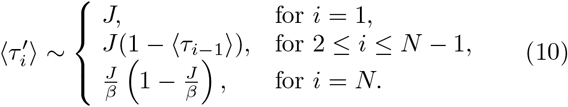

Using the asymptotics of 〈*τ_i_*〉 in different phases [22], the resulting densities at the boundaries and far from the right boundary (〈*τ*_*N*−*j*_〉, 1 ≪ *j* ≪ *N*) can be computed, as summarized in Table I. The asymptotics far from the left boundary (〈*τ*_*j*_〉, 1 ≪ *j* ≪ *N*) can be derived using the “particle-hole symmetry” [22]

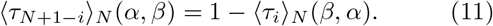

**TABLE I.**
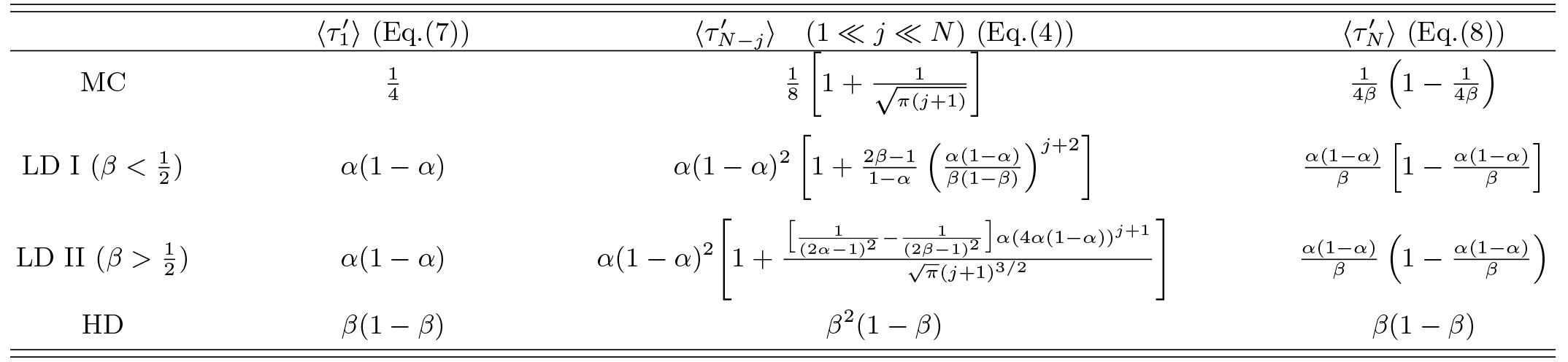
Asymptotics of *τ′* in the different phases of the classical 1-TASEP. These are obtained by combining equations (4), (7) and (8) with asymptotics given in [22]. The asymptotics far from the left boundary (〈*τ*_*j*_〉, 1 ≪ *j* ≪ *N*) can be derived using the “particle-hole symmetry” (11).

The fraction of isolated particles 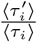 is given by

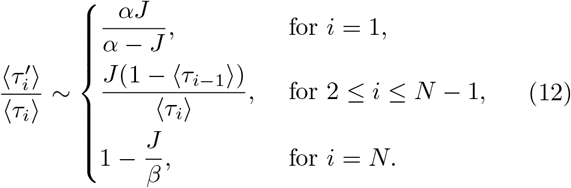

As shown in Fig. 2, there is good agreement between our asymptotic formulas and the exact results obtained from using the exactfrom using the exact (*τ_i_*)*_N_* in equations (4)–(8). We observed some large boundary effects, as the density of isolated particles at the boundaries is always larger than in the bulk. In the LD I regime 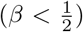, slow termination creates queuing so that the density of isolated particles decreases close to the end, in contrast to the total density. In the HD regime, high density creates a lot of stacked particles so the proportion of isolated particles is very small. In the MC regime, stacked particles are present more in the beginning of the lattice. As a result, the density of isolated particles in the bulk increases along the in the middle of the lattice. The apparent discontinuity in the middle of the lattice is due to the fact that we respectively employed in left and right parts of the lattice the asymptotics of the densities far from left and far from right of the boundaries, obtained from Table I. The resulting order of magnitude of the discontinuity gap in the middle is 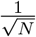, and it thus vanishes as the lattice length *N* increases. Interestingly, this gap can also be reduced to any arbitrary size by considering larger order approximations of the exact formulas for the densities found in [22]. Close to the boundaries, the formulas we used in Fig. 2 also lead to a mismatch with Monte Carlo simulations, which can be attributed to using asymptotics for positions far from the boundaries. This mismatch can be easily corrected by using the exact formulas for the densities at positions *N* − 1 and 2 (Equation 77 in [22]).

**FIG. 2.**
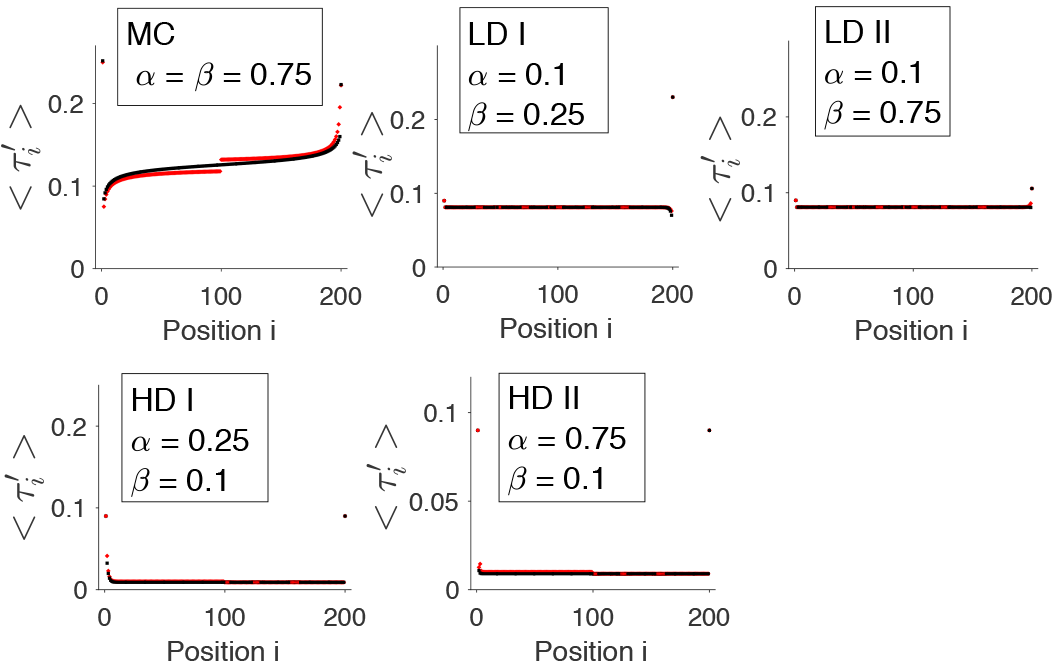
The density of isolated particles in different regions of the TASEP phase diagram. For the different regimes of the TASEP (see also Fig. 1), the asymptotic formulas from Table I (red points) are compared with the exact densities (black points) of isolated particles given by (4)–(8).

### C. The *ℓ*-TASEP with extended particles

During translation, ribosomes move along mRNAs by decoding one codon at a time, but occupy an extended space of ~ 10 codons. For that reason, it is also of interest to generalize our theoretical results to a process where particles occupy a certain size *ℓ* ≥ 1 (this process is usually called the *ℓ*-TASEP [26]). In this general case, using a matrix product to represent the steady-state solution leads to equations that are more complex, making the method employed above inapplicable (see Discussion). To cope with this complexity, we used a refined mean field approach introduced by Lakatos and Chou [2]. Although this approach cannot capture the variation of densities along the lattice as in the previous section, it well approximates the global average density and the current of particles. The key idea is to approximate the distribution of particles in the large-*N* limit by an equilibrium ensemble in which particles get uniformly distributed. Using such approximation, we obtained (Appendix D) that the density of isolated particles far from the boundaries, simply denoted 〈*τ′*〉, is given by

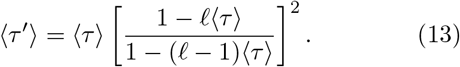

Using the asymptotic densities and currents found by Lakatos and Chou [2], we derived the asymptotics of 〈*τ′*〉. As for the *ℓ* = 1 case, the phase diagram can be decomposed into three parts (MC, HD, LD), separated by critical values 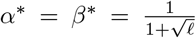. For *ℓ* = 1, we have 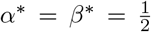, in agreement with the previous section. Combining (13) with the asymptotic density 〈*τ*〉 in the large-*N* limit [2], we obtained the following density of isolated particles in the bulk:

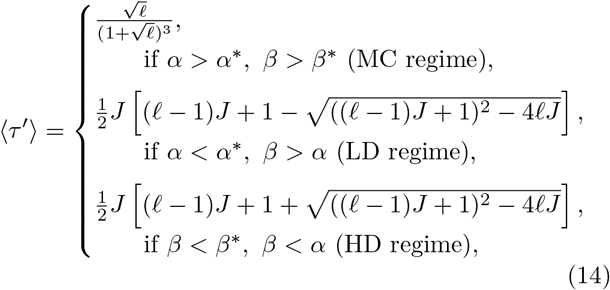

where *J* is the particle flux given by [2]

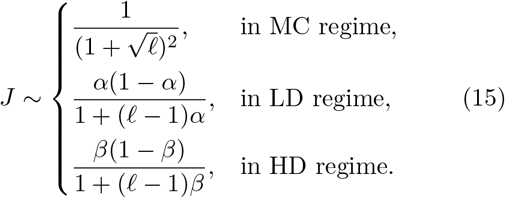

Near the entrance and exit, particles potentially get stacked on one side only. At the entrance, the density of isolated particles is, for *i* < *ℓ*,

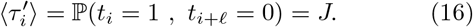

Hence, 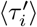 at the entrance is exactly given by the current flux *J*, as in the case of *ℓ* = 1. Near the exit, for *i* > *N* −*ℓ*, 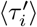 satisfies

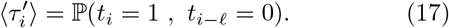

Using ℙ(*A* ⋂ *B*) = 1 − ℙ(*A^c^*) − ℙ(*B^c^*) + ℙ(*A^c^* ⋂ *B^c^*) and ℙ(*t*_*i*−*ℓ*_ = 1, *t*_*i*_ = 0) = *J* yields

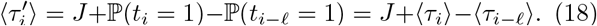

As the flux satisfies *J* = *β*〈*τ*_*N*_〉) = (*τ*_*N*−1_) = ··· = 〈*τ*_*N*−*ℓ*+1_), we obtained

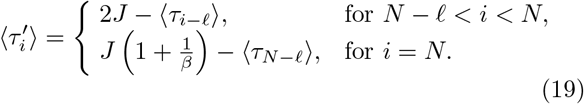

### D. Comparison with Monte Carlo simulations and estimation of interference rate

Combining (14), (16) and (19) leads to approximate densities of isolated particles along the lattice in the *ℓ*- TASEP. The isolated particle densities in the bulk (14) and near the entrance (16) depend only on the flux *J*, whereas near the exit the result (19) also depends on the density of particles located *ℓ* sites behind. In the LD regime, this density can be approximated by the density in the bulk [2]. In the other regimes, the density varies near the boundary, so using this approximation might be inaccurate (see Fig. 3). As Fig. 4a shows, however, our theoretical results agree well with the empirical densities of isolated particles obtained from Monte Carlo simulations, for specific values of (*α, β*) in the LD, HD and MC regimes (for a lattice of length 300, typical of the mRNA sequences we studied next). Contrary to the matrix method for the classical 1-TASEP model, the refined mean field approximation does not capture the variation of isolated particle densities across the lattice. However, this variation is much smaller than that of the total density, especially in regions of high traffic. Thus, assuming the density of isolated particles to be constant turns out to yield a better match with simulated data than when the same is done for the total density.

**FIG. 3.**
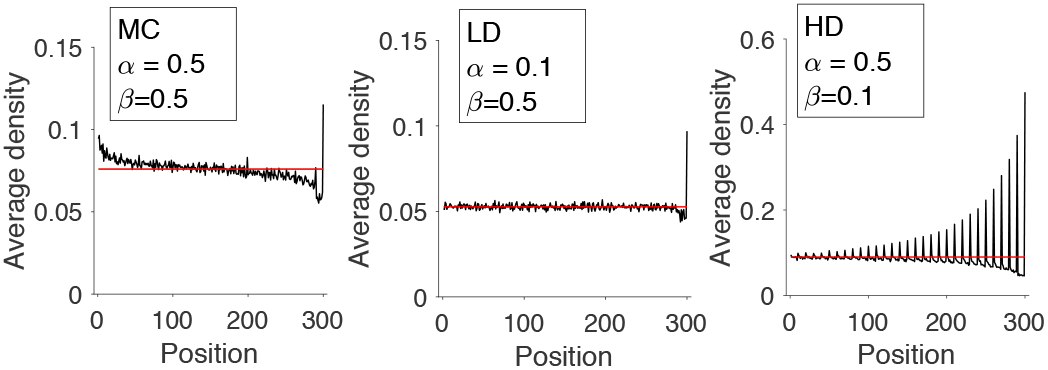
The density of particles in the *ℓ*-TASEP model. We simulated and plot (in black) the density of particles of the *ℓ*-TASEP (*ℓ* = 10) in the different regimes LD, HD and MC. In red, we plot the estimates of the density in the bulk from Lakatos and Chou [2].

More generally, we studied in Fig. 4b how the density and proportion of isolated particles vary as a function of *α*, for fixed values of *β*. Overall, our theoretical results were in good agreement with Monte Carlo simulations. Interestingly, whereas the total density (Fig. 4b) increase and reach a plateau after transitioning to the HD (when *β* < *β*\*) or the MC (when *β* > *β*\*) regime, the density of isolated particles follows a more complex pattern: First, there is a drop in the density of isolated particles when transition occurs from LD to HD. In contrast, we observed an increase in the total density, showing that most particles contributing to the density are stacked. Second, as *β* increases, the amplitude of the drop decreases until it becomes 0, when the MC regime replaces the HD regime. However, the maximum of 〈*τ′*〉 is not reached in the MC regime but in the LD regime before phase transition occurs. In other words, as the initiation rate increases, the level of interference increases faster than the global density. This was confirmed when we plotted the ratio 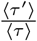 (Fig. 4b, right panels), showing a linear decrease from *α* = 0 to *α* = *β*, while the total density gets sublinear as *α* gets closer to *β*. The first-order Taylor expansion in *α* of 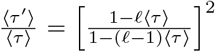 in the LD regime gives

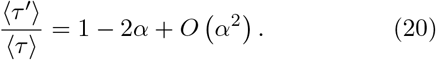

**FIG. 4.**
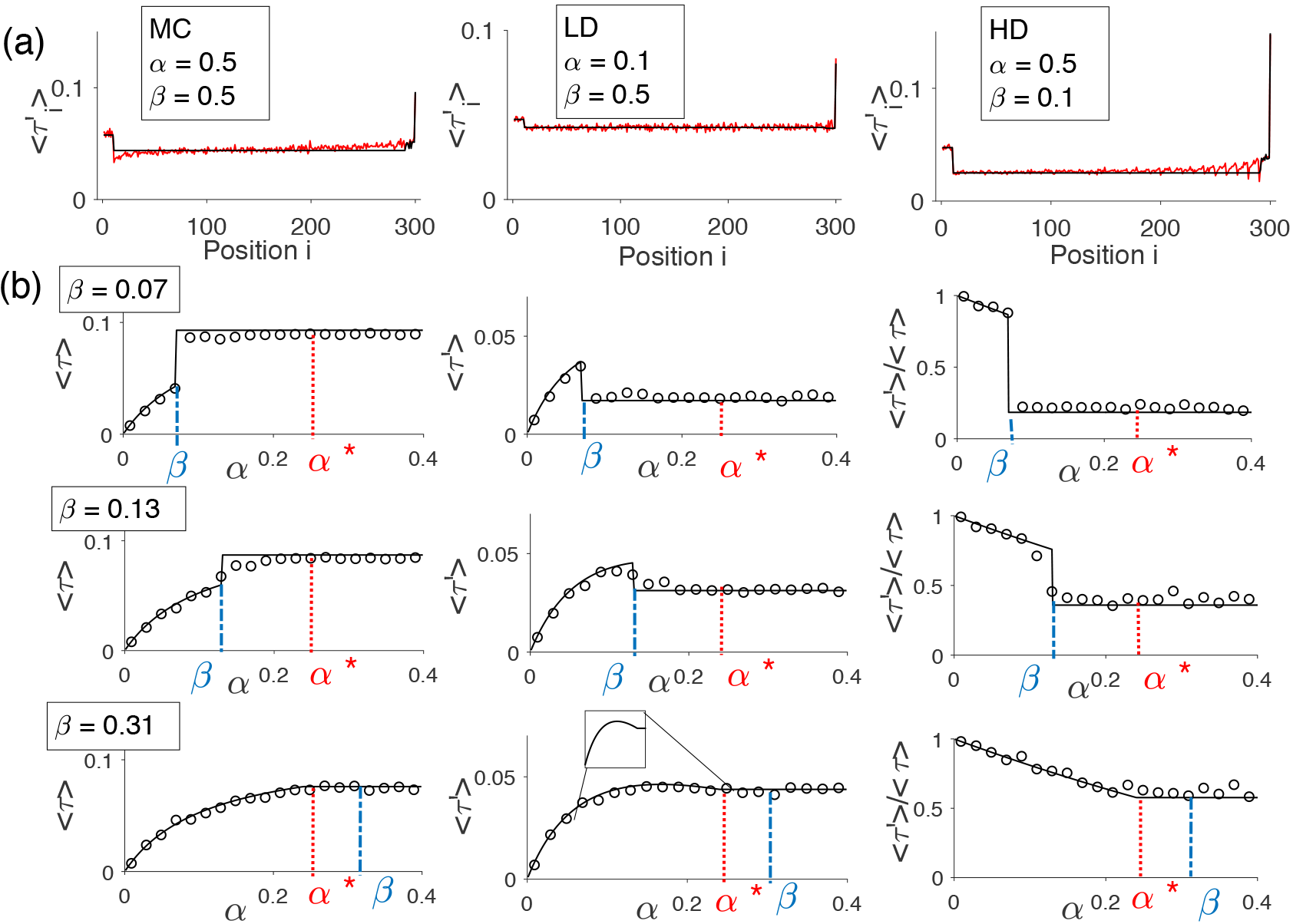
Comparison of the results from the reftned mean fteld approach with Monte Carlo simulations. **(a):** We simulated the TASEP with extended particles (size *ℓ* = 10, sample size = 10^9^) and plotted (in red) the densities of isolate particles in the three different regimes of the phase diagram. We compared these simulation results with the asymptotics obtain from (13), (16) and (19) (in black). **(b):** For fixed values of *β*, these plots show how the total density, the density of isolated particles, and their ratio vary as a function of *α*. The results obtained using Monte Carlo simulations (open circles) of the TASEP with extended particles (size of particles *ℓ* = 10, sample size of isolated particles 10^4^, lattice size = 400) are compared with the results obtained from the refined mean field approach (solid lines). Note that there are discontinuities when transitioning from LD to HD regime (first and second rows).

Interestingly, this formula does not depend on *ℓ* and using the formula obtained for the classical 1-TASEP model leads to the same result. To estimate the amount of interference associated with the dynamics of particles, we approximated the interference rate *I*, defined as the probability for a particle to get obstructed, as

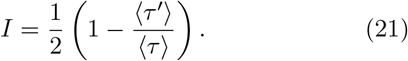

Using equation (20), we obtained that the interference rate is close to *α* in the LD regime.

### E. Generalization to larger isolation range

In the next section, one of our goals will be to determine whether stacked particles are detected in ribosome profiling experimental protocols. A problem is that we do not know *a priori* what is the exact range between two ribosomes that may prevent them from being detected. For this reason, we considered the density associated with isolation range *d*, denoted 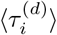, as

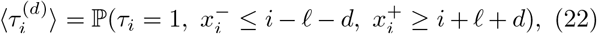

where 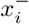 and 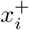 are the positions of the closest particles located before and after site *i*, respectively. In other words, 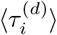 gives the steady-state density of particles under the *ℓ*-TASEP at position *i* such that the distance to their closest neighbor is at least *d + A*. In particular 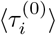 gives the total density of all particles, while 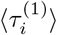 is equal to 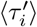, the density of isolated particles computed above. Following the same method as in the previous section, we obtained the following expression for particles in the bulk in the large-*N* limit:

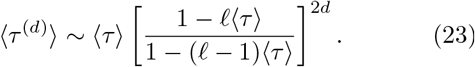

Hence, for two given isolation ranges *d* and *d′*, the associated fractions of isolated particles satisfy

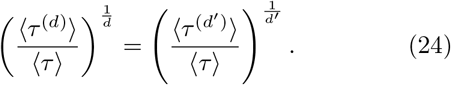

Therefore, we can generalize (21) to obtain a formula for the interference rate for an arbitrary isolation range *d* ≤ 1:

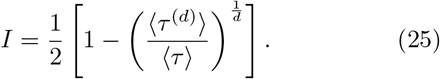

## APPLICATION

We applied our theoretical results to analyze ribosome profiling data and mRNA translation. Ribosomes are complex molecular machines (corresponding to particles in the TASEP) that move along mRNA (the lattice) to translate its associated sequence of codons into proteins. Once bound to the mRNA, ribosomes occupy a space of ~ 10 codons (*ℓ* = 10 in the TASEP). For the reader who is new to biology, more basics on how proteins are synthesized in a cell can be found in [4]. Briefly, the ribosome profiling procedure consists of using nuclease to digest translating ribosomes and get ribosome-protected mRNA fragments [15]. These fragments are then aligned to the mRNA sequence to produce a positional distribution of ribosomes along the mRNA. Assuming that there is no bias in ribosome detection and that sufficiently many fragments are observed, this distribution can be associated with the stationary average density of particles in the *ℓ*-TASEP. However, it is possible that the nuclease may fail to cleave stacked ribosomes [6, 17–19], so only the density of “isolated” ribosomes gets measured. Hence, the profile of ribosome counts along the mRNA produced by the experimental procedure might be different from the true profile (see Fig. 5). Whether the nuclease can cleave two nearby ribosomes is still in debate, as the digestion and its efficiency vary depending on the organism and the protocols which are used [27, 28].

**Fig. 5.**
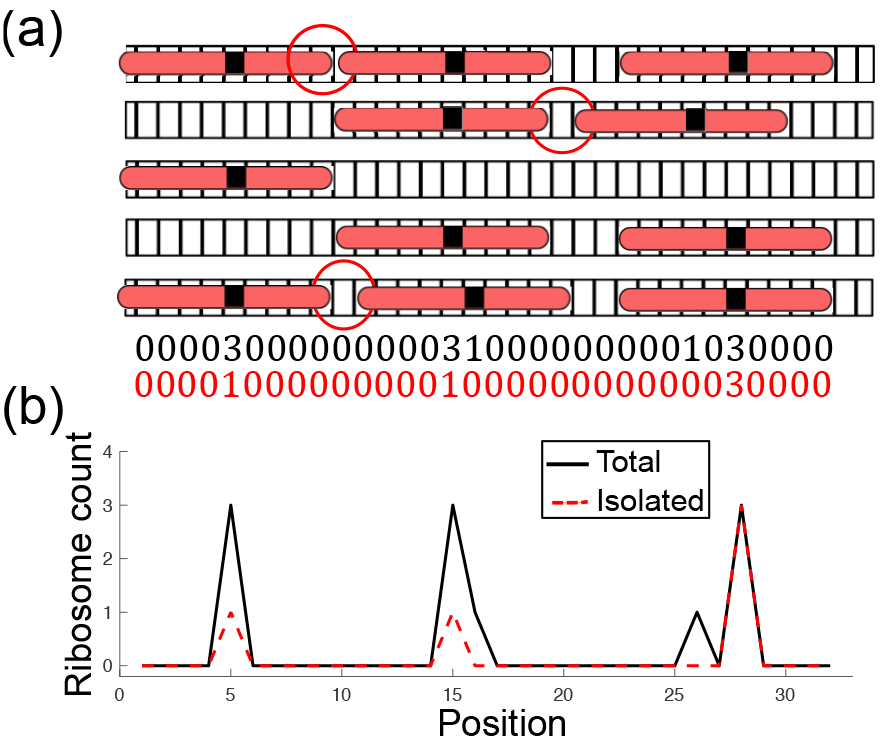
A schematic representation of ribosome proftling. **(a):** Positions of ribosomes along the mRNA are obtained by nuclease digestion and allow to count the number of ribosomes found at a specific position. However, it is possible that the nuclease cannot cleave stacked ribosomes [6, 14, 17–19]. **(b):** As a result, the profile of ribosome count along the mRNA recorded from isolated ribosomes (plotted in red) might be different from the true profile (plotted in black).

### F. Estimating the isolation range associated with non-detection of ribosomes

To assess the extent of non-detection of stacked ribosomes in an actual ribosome profiling dataset, we used publicly available data of *S. cerevisiae* from Weinberg *et al.* [29] (more details in Appendix E). The experimental protocol used for these data minimizes some of the other biases known to affect the ribosome profiling, such as sequence biases introduced during ribosome footprint library preparation and conversion to cDNA for subsequent sequencing, and mRNA-abundance measurement biases and other artifacts caused by poly(A) selection [29]. For a given gene, a measure of the average density of detected ribosomes is given by the so-called translation efficiency (*TE*) [23]. More precisely, the *TE* is given by the ratio of the RPKM measurement for ribosomal footprint to the RPKM measurement for mRNA, where RPKM denotes the number of Reads Per Kilobase of transcript per Million mapped reads. Hence, the *TE* is proportional to the average density of detected ribosomes per site of a single mRNA; in our notation, *TE* ∝ 〈*τ*^(*d*)^〉. To get the total density of ribosomes, we used another dataset from Arava *et al.* [30], obtained by polysome profiling, which is another technique giving, for a specific gene, the distribution of the number of ribosomes located on a single mRNA (and forming polysomes). While polysome profiling data is not biased by the possible omission of stacked ribosomes, the advantage of ribosome profiling is that it gives some local information about the ribosome occupancy.

Depending on the gap between two ribosomes that prevents them from being detected, the relation between the *TE* and the total average density *D* = 〈*τ*〉 is, according to equation (23),

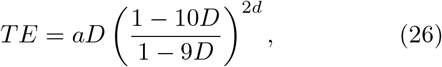

where *a* is the rescaling factor (specifically, *TE* = *a* · 〈*τ*^(*d*)^〉), that we estimate in practice in Fig. 6a, and *d* denotes the detection gap-threshold mentioned previously (if the gap between a ribosome and its closest neighbor is larger or equal to *d*, then it gets detected). Since a ribosome occupies 10 codons, the parameter *ℓ* in (23) is set to 10. In Fig. 6a, we plotted (26) for different values of *d* and compared it with the experimental data from Weinberg *et al.* and Arava *et al.* Our goal was then to determine which value of *d* leads to the best match with the experimental data. In Fig. 6b, we plotted the root mean square error between (26) and the experimental data, as a function of *d* and for the value of *a* corresponding to the linear fit to genes with total density less than 1 ribosome per 100 codons. We found that the minimum error is obtained when *d* is between 4 and 6. On the other hand, as *d* increases, the maximum value of *TE* that can be obtained using (26) decreases (Fig. 6a), potentially leading to some detected densities from experiment to be greater than the theoretical maximum of *TE*; we call such detected densities “anomalous” (as we shall see below, we can obtain a more refined estimate of the maximum possible detected density using an estimate of the termination rate *β* for each gene). In Fig. 6c, we plotted for each *d* the fraction of genes with anomalous detected densities. For *d* ≤ 3, no anomalous detected density was found, while the fraction becomes positive for *d* ≥ 4 (less than 1% for *d* = 4, ~ 2.5% for *d* = 6, and ~ 8% for *d* = 8). We concluded that the best values of *d* that both minimize the error and the fraction of anomalous detected density were obtained for *d* = 3 or 4. In agreement with our estimate, previous ribosome profiling experiments found disome fragments (accounting for the mapping of two ribosomes) of length ~ 65 nucleotides [19], suggesting that *d* = 3 (2 times 30 nucleotides plus 2 other codons).

**FIG. 6.**
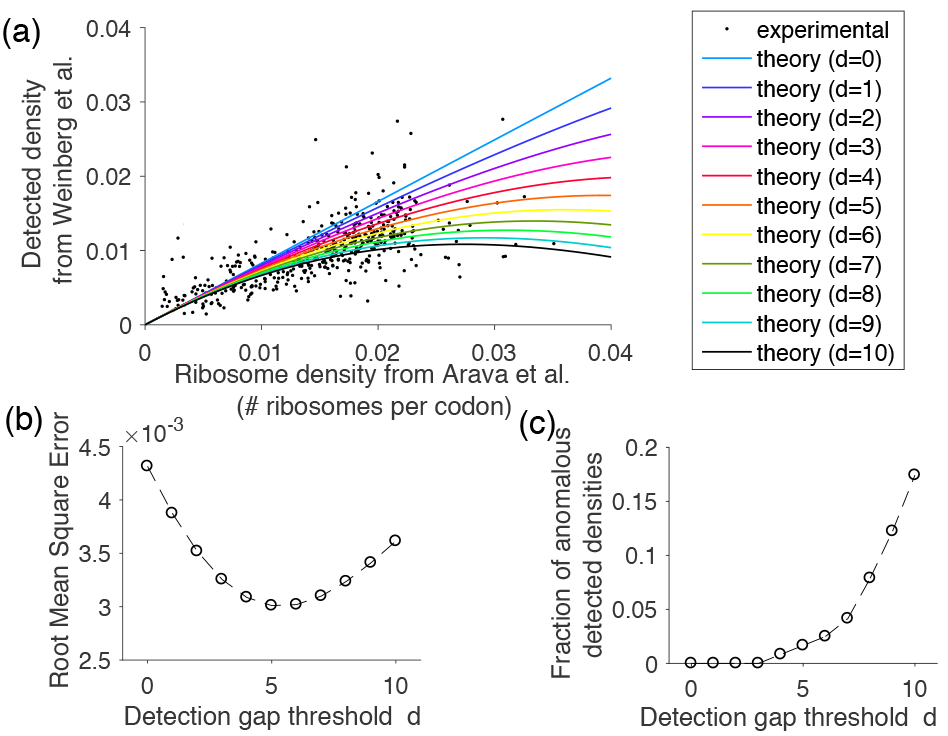
Estimation of undetected ribosomes from ribosome proftling experiment. **(a):** This plot shows experimental ribosome profiling data of *S. cerevisiae* from Weinberg *et al.* [29] against the total ribosome density obtained from polysome profiling by Arava *et al.* [30] (482 genes). Also shown are plots of 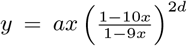, obtained from computing the density of detected particles of size *ℓ* = 10 as a function of the total density in the *ℓ*-TASEP (see (26)) with various isolation range *d* = 0,…, 10. We set *a* = 0.82, obtained by linear fit to genes with total density less than 1 ribosome per 100 codons. **(b):** For values of *d* ∈ {0,…, 10}, we plot the root mean square error obtained from comparing experimental data to the theoretical plots in **(a)**. **(c):** For *d* ∈{0,…, 10}, we plot the corresponding fraction of genes with anomalous detected densities, where a detected density said to be anomalous if it is larger than the theoretical maximum value implied by (26), used in **(a)**.

### G. Identiftability of initiation rates and flux from TE measurements

Under the *ℓ*-TASEP model in the LD regime, the *TE* is related (as shown in Fig. 7a) to the initiation rate *α* through equation (23) and the asymptotics of 〈*τ*〉 and *J* (given in [2]). Assuming that translation occurs in the LD regime (since translation is generally limited by initiation under realistic physiological conditions [31, 32]), we studied whether we could infer the gene-specific initiation rate *α* using our theoretical results. The detected density is bounded by ~ 0.02 ribosomes per codon in our dataset. From the plotted curves in Fig. 7a, this suggests that for *d* ≤ 5 and for all the experimental detected densities, there exists a value for the initiation rate satisfying (23). However, for *d* ≥ 3, the identifiability of *α* (i.e., the uniqueness of *α*) does not seem to be guaranteed.

**FIG. 7.**
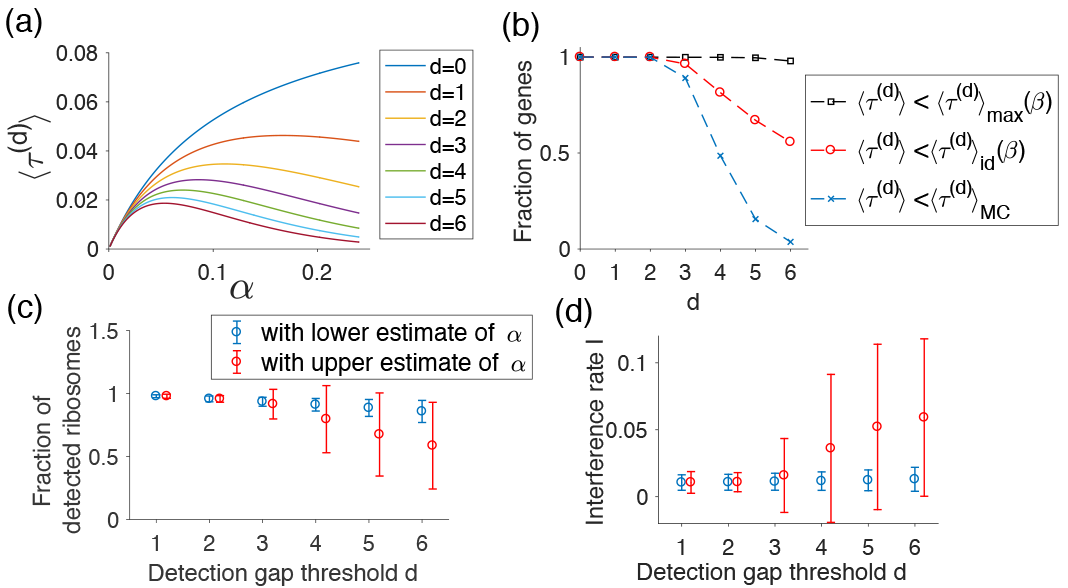
Analysis of initiation and interference. **(a):** For different values of isolation range *d*, we plot the density of isolated particles (see (23)) as a function of the initiation rate *α* in the LD regime. **(b):** For different ranges *d* of isolation, we studied the identifiability of the initiation rate. Black line: we estimated the fraction of genes for which there exists a corresponding value for the initiation rate *α*, such that the associated density of isolated particles is equal to the detected density. This happens when the detected density is less than 〈*τ*^(*d*)^〉_max_(*β*) (see (27)), where *β* is the inferred termination rate Red line: we estimated the fraction of genes for which the initiation rate can be inferred without ambiguity from the plotted curves in **(a)**, which happens when the detected density is less than 〈*τ*^(*d*)^〉_id_(*β*) (see (28)). **(c):** From our dataset of 3712 genes, we used (23) to estimate the fraction of detected ribosomes for different values of the detection gap threshold *d* ∈ {1,…, 6}. To compute these fractions when there is an ambiguity in identifying the initiation rate *α* (see **(b)**), we considered two possible estimates: a lower estimate and an upper one (see also Fig. 8a). The left plot represents the average fraction of detected ribosomes, with error bars indicating the standard deviation, using lower estimates (in blue) and upper estimates (in red) of *α*. **(d):** The same as in **(c)** for interference rates, using (25) (see also Fig. 8b).

More precisely, for a given gene and isolation range *d*, the theoretical maximal value of the *TE*, denoted 〈*τ*^(*d*)^〉_max_, is determined by the termination rate *β*, as

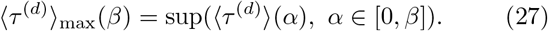

After estimating the termination rates from our ribosome profiling data (see Appendix F), in Fig. 7b we computed for different values of *d* the fraction of genes satisfying *T E′* ≤ 〈*τ*^(*d*)^〉_max_, where *T E′* is the *TE* normalized by the scaling factor *a* (see (26)). We found that all the genes satisfied this condition for *d* ≤ 5, before observing a small decrease for *d* = 6 (98%).

We further looked at the fraction of genes for which we can identify a unique initiation rate that matches the associated detected density with the measured *TE*. As *α* increases to its critical value min(*β, β*\*) (leading to a transition from LD to the other regimes), the density of isolated particles either only increases, or increases then decreases, to 〈*τ*^(*d*)^〉_id_(*β*), is determined by the termination rate *β*, given by

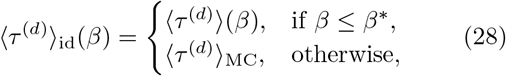

where 〈*τ*^(*d*)^〉_MC_, is determined by the termination rate *β* is the density of isolated particles in the MC regime. As a consequence, there is only one identifiable initiation rate in the LD region when *T E′* < 〈*τ*^(*d*)^〉_id_(*β*), and two when 〈*τ*^(*d*)^〉_id_(*β*) ≤ *T E′* ≤ 〈*τ*^(*d*)^〉_max_. In Fig. 7b, we computed the fraction of genes satisfying *TE* ≤ 〈*τ*^(*d*)^〉_id_. We found that all genes were then strictly identifiable for *d* ≤ 2, before the fraction starts to decrease for *d* = 3 (96%). For *d* ≥ 4, a significant fraction of genes (at least 19%) is not strictly identifiable. Thus, in the range of *d* associated with non-detection found from Fig. 6, the TE measurement may lead to some ambiguity in the initiation rates. In this case, two values of the initiation rate *α*_1_ < *α*_2_ lead to the same detected density: Although the total density for *α*_2_ is larger than for *α*_1_, there are also more closely stacked ribosomes that are not detected. Hence, the density of isolated particles is the same for both. As the flux is an increasing function of the initiation rate, such ambiguity also applies for inferring the flux.

### H. The fraction of detected ribosomes and interference rates

Upon estimating the threshold of gap distance between consecutive ribosomes leading to their non-detection and studying the identifiability of the initiation rate *α*, we then quantified the resulting fraction of detected ribosomes and the associated interference rate. As discussed above, for some values of *d* and 〈*τ*^(*d*)^〉, there may be two distinct values of *α*, and hence two distinct values of the total average density 〈*τ*〉, corresponding to the same 〈*τ*^(*d*)^〉. This implies that the fraction 〈*τ*^(*d*)^〉/〈*τ*〉 of detected ribosomes and the interference rate may not be uniquely determined for some values of *d* and 〈*τ*^(*d*)^〉. Indeed, for some of the experimentally observed *TE* values from Weinberg *et al.* [29], we encountered ambiguity in estimating *α* when *d* ≥ 3 (see Fig. 7b). Thus, when such ambiguity occurred, we considered both lower and upper estimates of *α*, and found their respective resulting fractions of detected ribosomes 〈*τ*^(*d*)^〉/〈*τ*〉 and interference rates (Fig. 7c and d). We obtained that for *d* = 3 or 4, suggested by Fig. 6b and c, the lower estimates of *α* lead to fractions of detected ribosomes lying between 91.2±5% and 93.5±3.5%. The upper estimates of *α* lead to smaller mean and larger variability (between 80±26% and 91.6±11.7%). As expected, we observed no substantial difference between the lower and upper estimates for *d* = 1 or 2 (since no gene presents any ambiguity). As *d* increases, however, the fraction of detected ribosomes decreases (notably because of the increasing fraction of genes with ambiguity). Interestingly, in contrast to these important variations, we observed that the interference rates corresponding to the lower estimates of *α* remain stable around 1% for all *d*, with only a slight increase of standard deviation from 0.5 to 0.9%. Somewhat larger variation is observed for the interference rates corresponding to the upper estimates of *α*, with ranges 1.5 ± 2.7% and 3.6 ± 5.5% for *d* = 3 and 4, respectively.

This difference in the amplitude between the fraction of detected ribosomes and interference rate can be explained theoretically, as illustrated in Fig. 8. When plotting the fraction 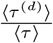 of detected ribosomes as a function of 〈*τ*^(*d*)^〉 (Fig. 8a), we observed that for large values of the fraction (associated with low *α*), the curves for different values of *d* were well separated, such that for 〈*τ*^(*d*)^〉 ~ 0.01 (corresponding to the range of our dataset), the fraction of detected ribosomes can vary between 98% (for *d* = 1) and 85% (for *d* = 6). In contrast, the interference rate takes approximately the same value for all *d* (~1%, see Fig. 8b). More generally, the formula (25) for interference rate shows that, as *d* increases, any observed decrease in the fraction 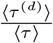 is compensated by the power 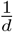. Furthermore, as *d* increases, the range of the ratio 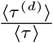 also increases (from 60 ~ 100% for *d* = 0 to 3 ~ 100% for *d* = 6), leading to larger differences between the lower and upper estimates, and higher variability across genes. In contrast, the interference rate remains bounded (by ~ 0.2), explaining its smaller variation across our dataset and different values of *d*.

**FIG. 8.**
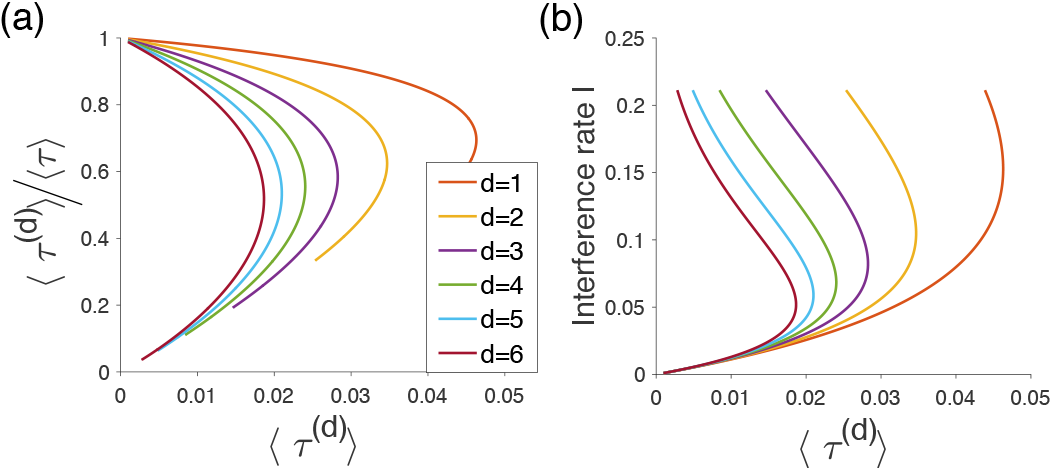
The fraction of isolated particles and interference rate as a function of 〈*τ*^(*d*)^〉. **(a):** For different isolation ranges *d* ∈ {1,…, 6}, we plot the fraction of isolated particles as a function of the average density of isolated particles 〈*τ*^(*d*)^〉, according to (23). Note that for given *d*, some values of 〈*τ*^(*d*)^〉 can lead to two possible fractions of isolated particles. **(b):** As in **(a)**, we plot the isolation rate as a function of the average density of isolated particles 〈*τ*^(*d*)^〉, according to (25). Note that for 〈*τ*^(*d*)^〉 ≤ 0.02 and all *d*, the initiation rates associated with the lower branch are very close.

## DISCUSSION

### I. Comparison with existing literature

In this article, we provided a complete analysis of the distribution of isolated particles in the TASEP model with open boundaries. This study was motivated by the possible non-detection of stacked ribosome in ribosome profiling, which is a recent experimental technique [23]. As shown in (3), the density of isolated particle is related to two- and three-point correlators, while most past analyses focused on computing the total density profile and the flux, which involve one- and two-point correlators. In the classical form of the model, we obtained exact analytic solutions using the matrix formulation originally developed by Derrida *et al.* [22]. We also obtained accurate asymptotic formulas in the limit of large *N* for different regimes of the phase diagram. In the past, the classical 1-TASEP has been studied in various geometric settings [20, 21], such as rings [33, 34] and networks [35, 36], or with more complex dynamics associated with pausing [37, 38], random rates [33, 39, 40] or multiple species [22, 34, 40–42], to name a few. A possible extension of our work would be to investigate the behavior of isolated particles in these different contexts. In many cases, the solution of the associated master equation can be found using a matrix formulation [20, 21, 40, 42], suggesting that the work presented here could be generalized.

We further studied the *ℓ*-TASEP model with extended particles of size *ℓ* and derived asymptotic formulas for densities using a refined mean field approach. In this more general case, the steady-state solution of the associated master equation can, in principle, also be written in the form of a generic matrix product [20, 43]. In practice, however, the associated algebra is rather complex, making it challenging to derive analytic results [2, 20]. To cope with this complexity, several approaches using mean field approximation have been developed [2, 4, 37, 44, 45]. Although the mean-field treatment may inaccurately capture the full profile in some regimes [37], it provides a more accurate approximation when the profile is restricted to isolated particles. More generally, the “level” of “mean-field” can also impact the quality of the approximation. At the simplest level, assuming a uniform distribution of particles without anti-correlations due to local interactions and using (13), one may obtain a rather poor approximation of the density of isolated particle, as the different regimes are not even separated correctly (all the cases in Fig. 4 would, for example, be considered as being in HD). Unlike this simple mean-field approach, the refined analytic approximation proposed by Lakatos and Chou [2] leads to formulas that show good agreement with simulations for current and bulk density [37]. In our work, we employed a similar approach to obtain a simple, accurate formula for the density of isolated particles with a given minimum distance to the closest neighbor. Higher “levels” of “mean-field” [44, 45] can help to improve the accuracy of the local density, but at the cost of losing analytic expressions, and possible existence of numerical instabilities and imprecisions [45].

The choice of lattice length (*n* = 300) in our comparison with Monte Carlo simulations was motivated by the typical size of the mRNA found in our dataset. As the length of the lattice increases, we expect the accuracy to improve, especially in the bulk, as the density would vary less. For much longer lattices, it would also be natural to study the hydrodynamic limit of the *ℓ*-TASEP with open boundaries. Interestingly, while previous studies derived a general PDE satisfied by the density for the *ℓ*-TASEP in the hydrodynamic limit [46, 47], a rigorous derivation, notably including that of boundary conditions, and analysis of the PDE to determine the associated phase diagram are still missing. We are currently exploring this research direction.

### J. Application to ribosome proftling data and comparison with other approaches

We applied our theoretical results to study mRNA translation using ribosome profiling data. In particular, our analysis suggests that the representation of the ribosome density may be biased by the non-detection of ribosomes with gap distance less than ~ 3 codons. In general, different protocols applied to different organisms can affect the nuclease action and in particular its ability to cleave ribosomes [28]. Hence, it would be interesting to apply our method to other datasets and other organisms to find possible differences in the detection gap distance. In particular, such differences could be visible near the terminal end of the transcript sequence, where slow termination can cause interference [48, 49]. In yeast (which is the organism studied in our dataset), no periodic peaks of density were detected in this region across multiple datasets [19, 50–57], suggesting non-detection of stacked ribosomes. In contrast, such peaks have been detected for other organisms and different protocols [58, 59].

Other methods have also been developed previously to infer the initiation rates associated with specific genes from polysome [9] or ribosome profiling [12]. These approaches used Monte Carlo simulations that can be computationally expensive. Using our theoretical results, it is possible to infer the initiation rate directly from the observed average detected density. Interestingly, we found that for our typical detection gap distance, some initiation rates were not uniquely identifiable (i.e., two initiation rates can lead to the same observed *TE* arising from isolated ribosomes), as having a higher initiation rate also creates higher interference that decreases the detected density. As a result, our work suggests that, for some genes, there could be ambiguity in identifying the initiation rate and the flux from *TE*, although this measurement has been widely used as a proxy for protein production [23].

We also provided robust estimates of the average rate of interference that ribosomes experience during translation. These estimates implicitly depend on the initiation rate and homogeneous elongation rate, but do not include other possible sources of interference due to local heterogeneities. More precisely, there is evidence of variation of the elongation rate along the transcript, especially in the first ~ 200 codons, leading to the so-called “5’ translational ramp” [23] (in another study [14], we quantified the extent of the interference created by this ramp). However, it has been shown that the average elongation speed along the transcript sequence is approximately constant around 5.6 codon/s [15], allowing the use of the homogeneous TASEP model as a first approximation of the translation dynamics.

Overall, our work shows how studying the interaction range of particles in exclusion process can help to get a better understanding of the process, and that it can be applied to problems where the data available are biased against this range. Similarly, while we focused here on isolated particles, our methods can be applied to situations where only aggregated particles following a transport process get detected.

## K. ACKNOWLEDGMENTS

This research is supported in part by a Math+X Research Grant from the Simons Foundation, and a Packard Fellowship for Science and Engineering. YSS is a Chan Zuckerberg Biohub investigator.

## Appendix A: Equations satisfted by the correlators in the TASEP

Averaging the master equation associated with the TASEP, the particle densities satisfy the following relations [60]:

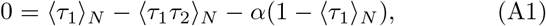

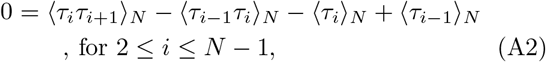

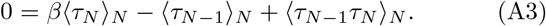

Note that (A2) implies 〈*τ_i_*(1 − *τ*_*i*+1_)〉_*N*_ = 〈*τ*_*i* − 1_(1 − *τ*_*i*_)〉_*N*_ for all *i* = 2,…, *N* − 1. This translation-invariant quantity is called the current (or flux) and is denoted by *J*. One can also relate the two-point correlators with the three-point correlators as

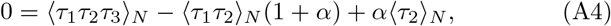

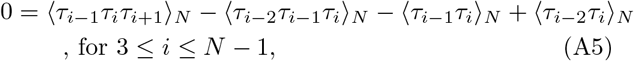

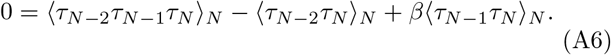

## Appendix B: Description of the matrix Ansatz used in the simple TASEP

To derive analytical expressions for the average densities of the TASEP, Derrida *et al.* [22] showed that the steady state probability of a given configuration can be derived using a matrix formulation as

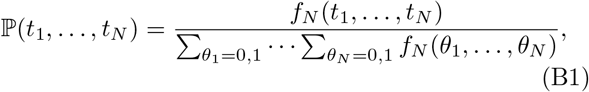

where

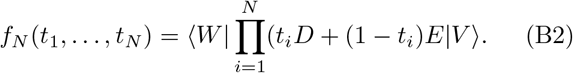

Here, *D* and *E* are infinite dimensional square matrices and |*V*〉 and 〈*W*| are column and row vectors respectively satisfying

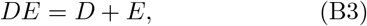

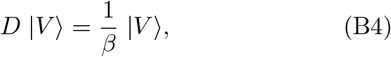

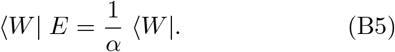

Using this formulation, the particle density can be derived as

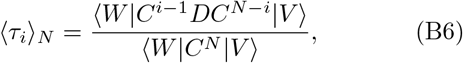

where *C* = *D* + *E*. More generally, for any given index set *i*_1_, *i*_2_,…, *i_k_* such that 1 ≤ *i*_1_ < ··· < *i_k_* ≤ *N*, we get

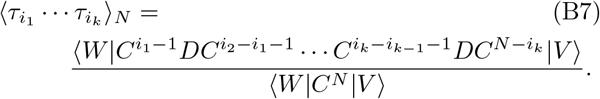

## Appendix C: Computing the density of isolated particles

Using the matrix Ansatz, we derive here an analytical expression for the average density of isolated particles 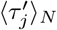. Our goal is to get 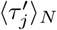 as a function of the average densities 〈*τ*_*j*_〉_*N*_. The density of isolated particles inside the lattice (2 ≤ *i* ≤ *N* − 1) is given by (see equation (3))

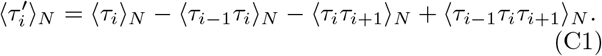

For 2 ≤ *j* ≤ *N* − 1, we first derive the expression of the two point correlators 〈*τ*_*j*_*τ*_*j*+1_〉*_N_* by summing equation (A2) over *i* ∈ {2,…, *j*} and using the boundary equation

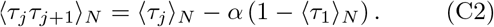

Similarly, for 3 ≤ *j* ≤ *N* − 1, summing equation (A5) from *i* = 3 to *j* and using boundary equations (A1) and (A4) gives

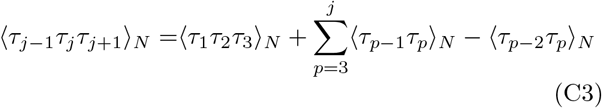

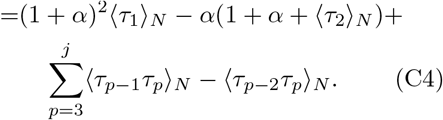

Using the matrix formulation and the identities *DCD* = *D*(*DC* − *DE* + *ED*) = *DDC* − *DC* + *CD*, we get

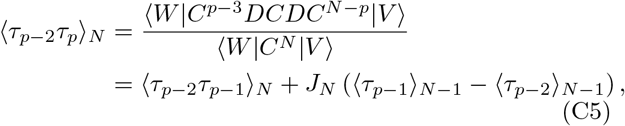

where 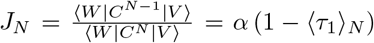 is the particle current at steady state [22]. Combining (C5) with (C4) and using (C1) and (A1) yield the result for the threepoint correlator

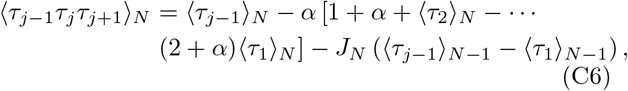

for 3 ≤ *j* ≤ *N* − 1. Using (A1) and (A4), this equation is also true for *j* = 2. Using (C6), (C2) and (3) gives us the formula for the density of isolated particles, for 2 ≤ *i* ≤ *N* − 1

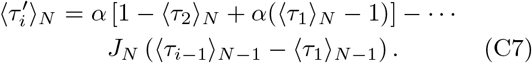

Finally we can use *J*_*N*_ = *α* (1 − 〈*τ*_1_〉_*N*_) to write the above formula in a more compact notation, as

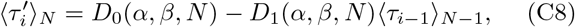

where

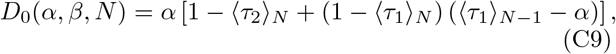

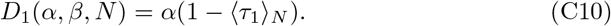

Similarly, using equations (A1) and (A3) at the boundaries yields

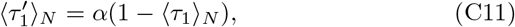

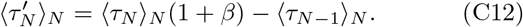

## Appendix D: Density of isolated particles in the bulk for the *ℓ*-TASEP

We compute here an estimate of the density of isolated particles of size *ℓ* in the bulk (〈*τ_i_*〉, 1 ≪ *i* ≪ *N* − *ℓ*). To do so, we use an approximation from Lakatos and Chou [2], assuming that the number of states of *n* particles of length *ℓ*, confined to a length of *N′* ≥ *nℓ* lattice sites, is given by the partition function [61]

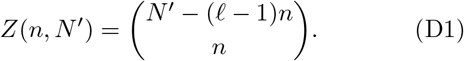

For a given position *i* ∈ {1,…, ≤ *N* − *ℓ*, we introduce 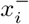 and 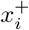 as the positions of the closest particles to the left and the right of *i*, respectively, so we get

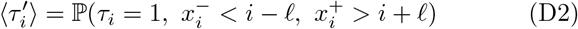

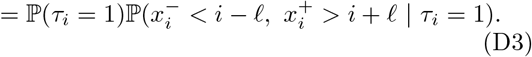

Assuming 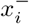 and 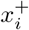 being independent ieilds

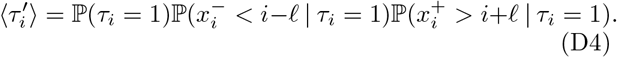

Using (D1), the probability 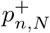 that 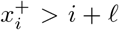, conditioned on *τ_i_* = 1 and there being *n* particles in the window [*i* + *ℓ*: *i* + *ℓ* + *N′* − 1] is

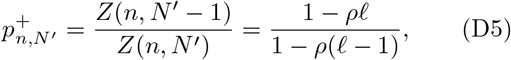

where 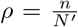. When *n* and *N′* get large and assuming the density of particles in the bulk of the lattice to be approximately constant (denoted 〈*τ*〉, we can replace 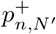 in equation (D5) by 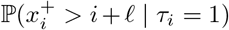 and 〈*τ*〉, respectively, which gives

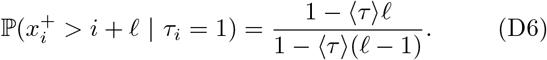

Similarly, we obtain 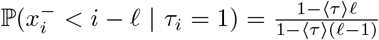 Combining these relations and replacing ℙ(*τ_i_* = 1) by 〈*τ*〉 in equation (D3), we obtain that the density of isolated particles in the bulk, simply denoted 〈*τ′*〉, is given by

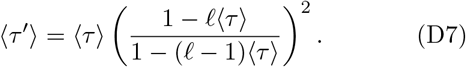

Similarly, for isolation range *d*, we obtain

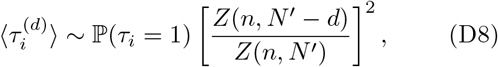

which simplifies to the following expression in the large-*N* limit:

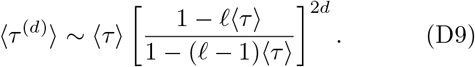

## Appendix E: Experimental dataset

The flash-freeze ribosome profiling data from Weinberg *et al.* [29] can be accessed from the Gene Expression Omnibus (GEO) database with the accession number GSE75897. To map the A-sites from the raw short-read data, we used the following procedure: We selected the reads of lengths 28, 29 and 30 nt, and, for each read, we looked at its first nucleotide and determined how shifted (0, +1, or 1) it was from the closest codon’s first nucleotide. For the reads of length 28, we assigned the A-site to the codon located at position 15 for shift equal to +1, at position 16 for shift equal to 0, and removed the ones with shift 1 from our dataset, since there is ambiguity as to which codon to select. For the reads of length 29, we assigned the A-site to the codon located at position 16 for shift equal to +0, and removed the rest. For the reads of length 30, we assigned the A-site to the codon located at position 16 for shift equal to 0, at position 17 for shift equal to −1, and removed the reads with shift +1.

## Appendix F: Estimation of termination rates

For a given profile (*P*_1_,…, *P_N_*) containing the number of footprints with A-site detected at each position, we estimate the associated scaled termination rate as

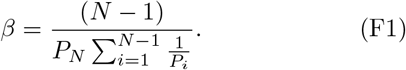

Such estimation is valid when there is little ribosomal interference, such that the elongation rate can be approximated by the inverse of the profile [14]. In another study [14], we developed a more refined inference procedure that uses these rates as first estimates (this method applies for genes with high footprint coverage), leading to excellent agreement between the observed and simulated profiles for the same dataset used here. As in average, our refined procedure lead to correction for ~ 1.57 site per gene, these “naive” estimates are valid over a large majority of the sites.

